# Multiple zebrafish *atoh1* genes specify a diversity of neuronal types in the zebrafish cerebellum

**DOI:** 10.1101/098012

**Authors:** Chelsea U. Kidwell, Chen-Ying Su, Masahiko Hibi, Cecilia B. Moens

## Abstract

The basic Helix-Loop-Helix transcription factor Atoh1 is required for the specification of multiple neuron types in the mammalian hindbrain including tegmental, precerebellar output neurons and cerebellar granule neurons. How a single proneural gene specifies so many neuron types from a single progenitor zone, the upper rhombic lip (URL), is not known. Here we use the zebrafish to explore the role of *atoh1* in cerebellar neurogenesis. Using transgenic reporters we show that zebrafish *atoh1c*-expressing cells give rise to tegmental and granule cell populations that, together with previously described *atoh1a*-derived neuron populations, resemble the diversity of *atoh1* derivatives observed in mammals. Using genetic mutants we find that of the three zebrafish *atoh1* paralogs, *atoh1c* and *atoh1a* are required for the full complement of granule neurons in the zebrafish cerebellum. Interestingly, *atoh1a* and *atoh1c* specify non-overlapping granule populations, indicating that fish use multiple *atoh1* genes to generate granule neuron diversity that is not detected in mammals. By live imaging of neurogenesis at the URL we show that *atoh1c* is not required for the specification of granule neuron progenitors but promotes their delamination from the URL epithelium and this process is critical for neuronal maturation. This study thus provides a better understanding of how proneural transcription factors regulate neurogenesis in the vertebrate cerebellum.

**Summary statement:** *atoh1* genes specify distinct populations of tegmental and granule neurons in the zebrafish hindbrain and promote their delamination from the neuroepithelium, a process critical for neuronal maturation.

## INTRODUCTION

The cerebellum is well known for its importance in motor coordination necessary for the generation of smooth and skillful movements (Leto et al., 2015). In all vertebrates, the cerebellum is composed of excitatory glutamatergic granule neurons and inhibitory GABAergic Purkinje neurons organized into a three-layered structure consisting of a superficial molecular layer, an underlying Purkinje layer, and a deep granule neuron layer (Butts et al., 2014; Hashimoto and Hibi, 2012; Leto et al., 2015). During mammalian development, granule neuron progenitors are born in the dorsal-most anterior hindbrain, a region called the upper rhombic lip (URL) while Purkinje neurons are born in the subjacent ventricular zone (Leto et al., 2015). Granule neuron progenitors delaminate from the URL epithelium and migrate tangentially from the URL across the surface of the presumptive cerebellum where they undergo transit-amplifying cell divisions before migrating radially to form the internal granule layer. Once in the internal granule layer, granule neurons elaborate axons that bifurcate in the molecular layer to form parallel fibers that synapse with Purkinje neurons (Altman and Bayer, 1997).

The origins and circuitry of cerebellar neurons is conserved in zebrafish with some differences (Kani et al., 2010). Zebrafish granule neurons migrate anteriorly from the URL to populate the Corpus Cerebelli (CCe), a structure homologous to the mammalian cerebellar vermis which is thought to control body positioning and into an anterior progenitor zone, the Valvula (Va). Granule neurons in fish also migrate laterally from the URL into the eminentia granularis (EG) and the caudal Lobus Caudalis (LCa), homologous to the mammalian flocculonodular lobe, to control motor coordination in response to vestibular information (See schematic in Fig. 1) (Kani et al., 2010; Kaslin et al., 2009; Matsui et al., 2014; Volkmann et al., 2010; Volkmann et al., 2008). While granule neuron progenitors have been observed to divide after delaminating from the URL (Kani et al., 2010), they do not undergo the massive amplification characteristic of the mammalian EGL (Butts et al., 2014; Chaplin et al., 2010).

**Figure 1.**
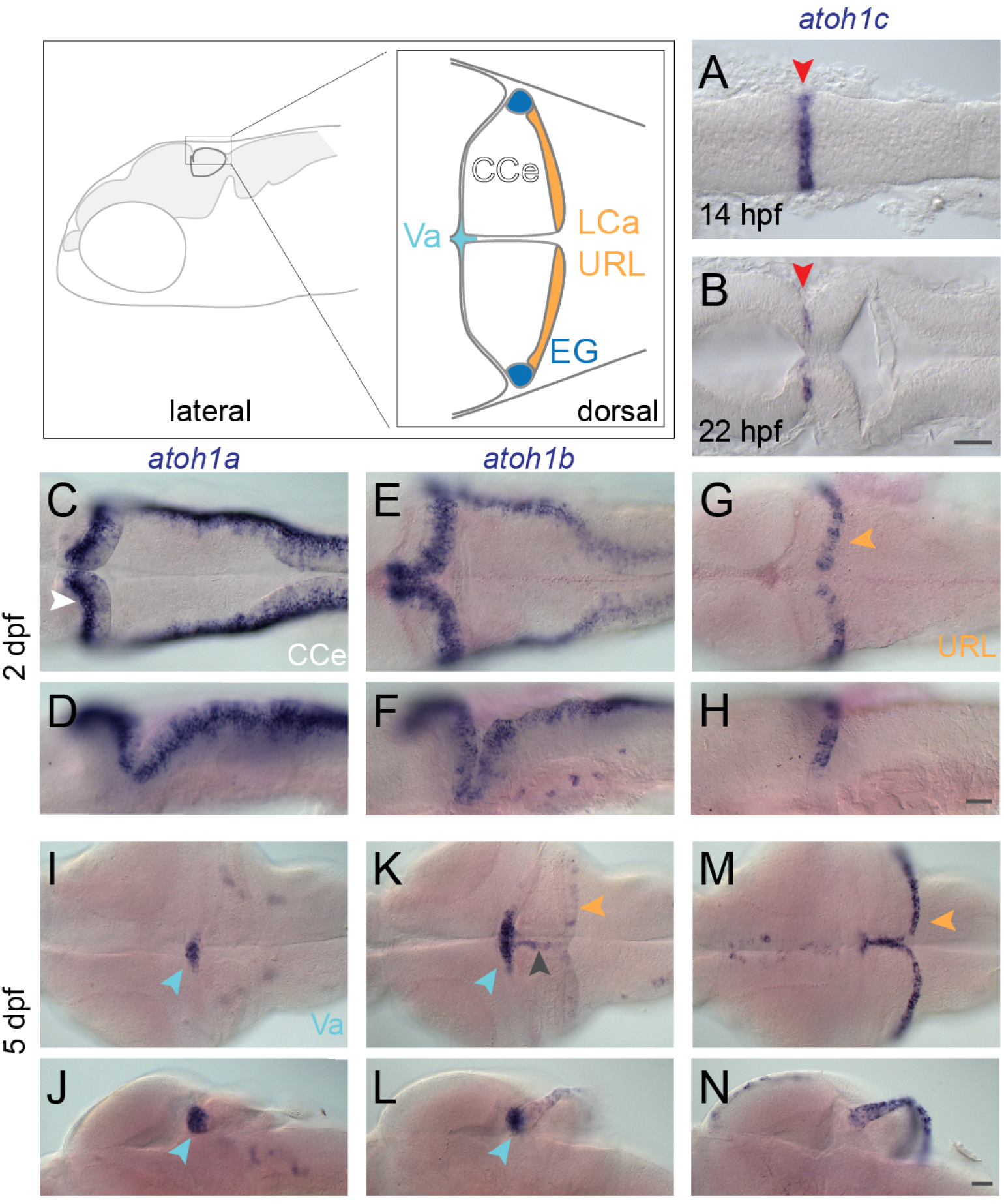
*atoh1* gene expression in the developing zebrafish cerebellum. Schematic depicts regions of the zebrafish cerebellum in lateral (left panel) and dorsal (right panel) views. Throughout the manuscript, white arrowheads indicate the CCe, light blue arrowheads indicate the Va, orange arrowhead indicate the URL (at early stages) or the LCa (at later stages) and dark blue arrowheads indicate the EG. Red arrowheads indicate the position of the MHB. A-N: RNA *in situ hybridization* in wild-type embryos with *atoh1a* (left column), *atoh1b* (middle column), and *atoh1c* (right column) at 14 hpf (A) 22 hpf (B), 2 dpf (C-H), and 5 dpf (I-N). Dorsal (A,B,C,E,G,I,K,M) or lateral (D,F,H,J,L,N) views are shown with anterior to the left. Gray arrowhead indicates midline of CCe. CCe, corpus cerebelli; EG, eminentia granularis; LCa, lobus caudalis cerebelli; URL, upper rhombic lip; Va, valvula cerebelli; Scale bars: 50 uM.

Atoh1, a highly conserved basic-Helix-Loop-Helix proneural transcription factor, has been shown to be vital for neuron specification in the hindbrain (Rose et al., 2009; Wang et al., 2005). Genetic lineage tracing in the mouse has demonstrated that *atoh1*-expressing progenitors in the URL give rise sequentially to diverse excitatory neuron types within the anterior hindbrain and cerebellum from 9.5 to 19 days post conception, including ˜9 tegmental neuron groups, deep cerebellar output neurons and all cerebellar granule neurons (Akazawa et al., 1995; Ben-Arie et al., 1997; Ben-Arie et al., 2000; Bermingham et al., 2001; Englund et al., 2006; Gray, 2008; Green et al., 2014; Machold and Fishell, 2005; Rose et al., 2009; Wang et al., 2005). How *atoh1* specifies so many neuronal types from a single progenitor zone is still unknown.

Zebrafish have three *atoh1* genes: *atoh1a, 1b*, and *1c*, which are expressed in overlapping but distinct progenitor domains within the rhombic lip. Initial studies have described the expression of the three zebrafish *atoh1* genes, and lineage tracing of atoh1a-expressing progenitors showed that they give rise to diverse neuronal cell types including tegmental neurons, cerebellar output neurons and granule neurons in the CCe and Va (Chaplin et al., 2010; Kani et al., 2010). The absence of *atoh1a*-derived granule neurons in the EG and LCa suggested that other *atoh1* genes may be responsible for granule neuron diversity in fish, however this has not been further explored.

We sought to discover the role of zebrafish *atoh1* genes in the generation of neuronal diversity in the cerebellum, and to take advantage of the live imaging possible in zebrafish to study how *atoh1* genes control cerebellar progenitor migration and differentiation at high spatial and temporal resolution. Using single and compound mutants we describe a predominant role for *atoh1c* and a lesser role for *atoh1a* in the specification of cerebellar granule neurons in zebrafish. Using long-lived *atoh1a* and *atoh1c* reporters to follow their derivatives through larval development, we find that *atoh1a* and *atoh1c* specify non-overlapping granule and tegmental populations, indicating that fish use multiple *atoh1* genes rather than using a single *atoh1* gene over an extended developmental time to generate neuronal diversity, and revealing an unexpected diversity within the granule neuron lineage. With the use of live imaging, we discovered that *atoh1c* expression at the rhombic lip is required not for cell cycle exit or initial granule neuron differentiation but for granule neuron progenitors to delaminate from the URL epithelium and initiate migration, a critical early event in neurogenesis.

## RESULTS

### Zebrafish *atoh1* genes are expressed sequentially at the URL

To determine the expression dynamics of the *atoh1* genes during embryonic cerebellar development, we carried out a time-series RNA *in situ* hybridization analysis.

Consistent with previous reports, we found that all three *atoh1* homologues are expressed in the URL beginning from 1 day post fertilization (dpf) and persist beyond 5 dpf (Fig. 1) (Chaplin et al., 2010; Kani et al., 2010). In addition, we identified an early transient *atoh1c* expression domain at the mid-hindbrain boundary (MHB), anterior to the presumptive URL. This expression domain is detected starting at 14 hpf until 30 hpf (Fig. 1A,B) at which point it is extinguished and expression in the URL begins at 2 dpf and persists until our analysis end point, 5 dpf (Fig. 1G,H,M,N). Both *atoh1a* and *atoh1b* are expressed in the URL beginning at 24 hpf (not shown), but by 5 dpf their expression is largely restricted to the Va, a progenitor zone in the anterior-most region of the cerebellum (Fig. 1C-F, I-L) (Kani et al., 2010; Kaslin et al., 2009). Although *atoh1b* is primarily expressed in the Va region at 5 dpf, weak expression in also detected in the upper rhombic lip and at the midline of CCe (Fig. 1K,L, gray and orange arrowheads) where *atoh1c* is strongly expressed (Fig. 1M,N).

### An early isthmic domain of *atoh1c* expression gives rise to tegmental neurons related to the Locus Coeruleus

The early expression domain of *atoh1c* lies at the MHB, an important signaling center required for early development of both the midbrain and hindbrain (Rhinn and Brand, 2001). Given the location of the early *atoh1c* expression domain we wondered whether MHB patterning network played a role in its regulation. The position of the MHB lies at the interface of Otx2 and Gbx2 and is reciprocally maintained by Wnt1 expression anterior to the boundary and Fgf8 expression posterior to the boundary.

Double RNA *in situ* hybridization with *otx2* and *gbx2*, demonstrated that *atoh1c* is expressed in a few cells immediately posterior to the boundary (Fig. S1A,B). Not surprisingly, MHB *atoh1c* expression is lost under conditions where FGF and Wnt signaling are blocked early in development by heat-inducible expression of dominant negative (dn) FGFR1 or dnTCFΔC, respectively (Fig. S1E-G) (Lee et al., 2005; Martin and Kimelman, 2012). To investigate the nature of this requirement, we made chimeras in which cells expressing dnFGFR1 or dnTCFΔC and a transgenic *atoh1c* reporter (see below) contributed sparsely to the MHB region. In these chimeras, dn-expressing cells are cell-autonomously blocked for FGF or Wnt signaling, but the MHB morphogenetic program occurs normally in the surrounding wild-type cells. Whereas non-dn-expressing cells at the MHB expressed the *atoh1c* reporter (n=10/10 chimeric embryos; Fig. S1H), cells expressing dnFGFR1 or TCFΔC at the MHB never expressed the *atoh1c* reporter (n=0/32 chimeric embryos; Fig. S1I,J). Thus, the tightly restricted expression of *atoh1c* to progenitors at the MHB is because cells must be able to perceive both Wnt and FGF signals in order to express *atoh1c.*

In order to visualize the atoh1c-derived cell populations *in vivo*, we generated an *atoh1c* transgenic reporter, *TgBAC(atoh1c:gal4FF)fh430* by BAC recombineering (Fig. 2A-L) (Bussmann and Schulte-Merker, 2011). When crossed to *Tg(UAS:kaede)s1999t* (Scott et al., 2007), our *TgBAC(atoh1c:gal4FF)fh430* driver (hereafter referred to as *Tg(atoh1c::kaede)* for simplicity) recapitulates all aspects of endogenous *atoh1c* expression at the MHB and URL (Fig. 2A-L). Taking advantage of the long-lived nature of the photoconvertible Kaede fluorescent protein (Ando et al., 2002; Caron et al., 2008) we were able to follow the fate of the MHB *atoh1c+* progenitor pool after *atoh1c* mRNA expression at the MHB was extinguished. Starting at 20 hpf, the MHB *Tg(atoh1c::kaede)*+ population migrates ventrocaudally to rhombomere 1 (r1) ventral to the presumptive cerebellum (the tegmentum) and gives rise to bilateral comma-shaped nuclei consisting of about 20 neurons by 2 dpf (Fig. 2A-F, O). Since the onset of *atoh1c* expression at the URL at 3 dpf complicates the subsequent lineage analysis of MHB-derived *atoh1c* neurons (see below for further discussion of *atoh1c* URL derivatives), we distinguished the MHB-derived *atoh1c* lineage by photoconverting the Kaede+ MHB domain at 22 hpf, before the onset of URL expression and found that the majority of neurons present in ventral r1 are Kaede Red+ indicating that they originated from the *atoh1c+* MHB progenitor domain (Fig. 2M-P).

**Figure 2.**
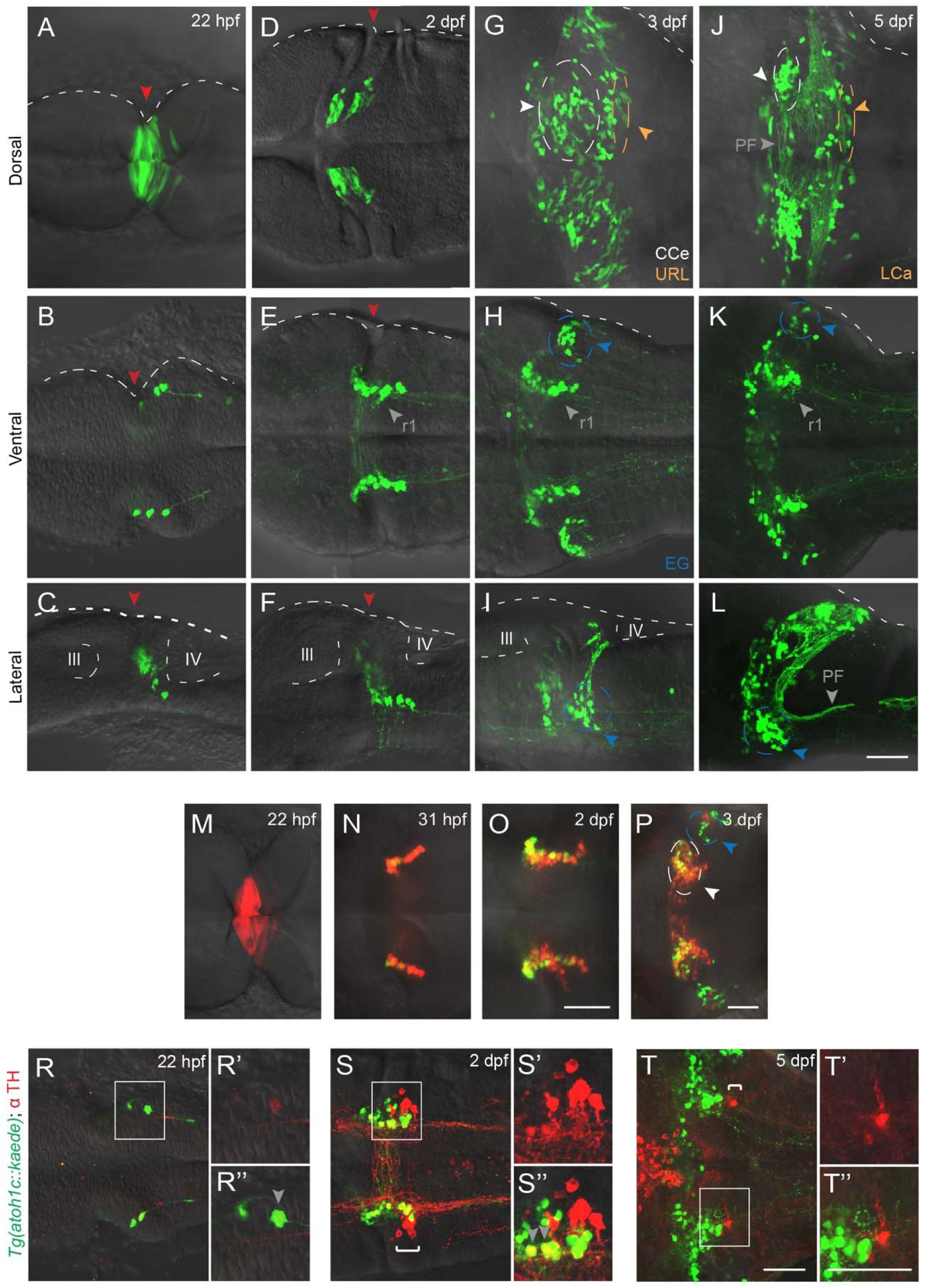
atoh1c-expressing progenitors give rise to to ventral r1 and cerebellar granule neurons. A-L: *Tg(atoh1c::kaede)* transgene expression in fixed embryos stained with anti-Kaede antibody in dorsal (A,D,G,J), ventral (B,E,H,K) and lateral (C,F,I,L) views at 22 hpf (A-C), 2 dpf (D-F), 2 dpf (G-I) and 5 dpf (J-L). Arrowheads follow the color code described in Fig. 1 legend or are labeled as follows: PF: parallel fibers, r1: MHB-derived neurons in ventral r1. M-P: live imaging after photoconversion of Kaede (green to red) of MHB *atoh1c+* cells at 22 hpf confirms that this *Tg(atoh1c::kaede)*+ progenitor domain gives rise to ventral r1 neurons. Dorsal (M) and ventral focal planes (N-P). R-T: *Tg(atoh1c::kaede)*+ cells (green) transiently express TH (red; gray arrowheads) from 22 hpf to 2 dpf and lie adjacent to the LC (indicated by white bracket). All images oriented with anterior to the left at time points as indicated. Scale bars: 50 μM.

The migratory path from the MHB of these *atoh1c+* neurons and their final position in r1 are highly reminiscent of the Locus Coeruleus (LC; (Chiu and Prober, 2013; Guo et al., 1999)). The LC is a noradrenergic neuronal population present in all vertebrates that controls arousal. At 5 dpf, the *Tg(atoh1c::kaede)+* neurons lie immediately anterior to the LC and have overlapping contralateral and longitudinal projections with the LC neurons (Fig. 2R-T) (Guo et al., 1999). Furthermore, the *Tg(atoh1c::kaede)*+ r1 neurons express the biosynthetic enzyme tyrosine hydroxylase (TH) during their migration (Fig. 2R, S, gray arrowheads), However, the *atoh1c+* r1 neurons are not the LC themselves, as they eventually turn off TH while the neurons of the LC maintain it (Fig. 2T). Given the similarities between these two populations, we hypothesize that the *Tg(atoh1c::kaede)+* r1 neurons may be functionally connected to the LC and be a part of the arousal circuit.

### Atoh1c is cell-autonomously required for the specification of multiple granule neuron populations in the zebrafish cerebellum

By 3 dpf, the majority of the *Tg(atoh1c::kaede)*+ cells in the cerebellum have migrated to regions populated by granule neurons: the CCe, EG and LCa (Fig. 2G-L, Movie 1). Their axons appear starting at 3.5 dpf and resemble granule neuron parallel fibers (Hibi and Shimizu, 2012; Takeuchi et al., 2015). In addition to the parallel fibers across the surface of the cerebellum (Fig. 2J, gray arrowhead), granule neurons of the EG and LCa also project parallel fibers out of the cerebellum to a cerebellar-like structure in the dorsal hindbrain (Fig. 2L, gray arrowhead) (Bae et al., 2009; Takeuchi et al., 2015). Given the location and projections of the *Tg(atoh1c::kaede)*+ population, we conclude that they are granule neurons of the CCe, EG, and LCa. atoh1c-derived granule neurons are not detected in the Va, where many atoh1a-derived granule neurons are located (Kani et al., 2010).

To directly assess the function of Atoh1c, we used with transcription activator-like effector nucleases (TALENs) to generate a 122 bp deletion (fh367) that results in a premature stop before the Atoh1c DNA binding domain. Expression of *cerebellin12* (*cbln12*), a marker of all mature granule neurons (Takeuchi et al., 2016), is strongly reduced in the CCe, EG and LCa in *atoh1c* mutants (Fig. 3A,B). Expression of the *vesicular glutamate transporter 1 (vglut1)* mRNA (Fig. 3C,D) and protein (not shown), which mark the glutamatergic neurons in the cerebellum, is similarly reduced. The majority of the *Tg(atoh1c::kaede)*+ cells accumulate at the URL in *atoh1c^fh367^* mutants, where they strongly express *atoh1c* mRNA (Fig. 3G compare to Fig. 1M,N), retain a neural progenitor-like morphology, and elaborate comparatively few parallel fibers (Fig. 3E,F compared to Fig. 2G,I,J,L; Movie 2). Thus *atoh1c* contributes to the specification of granule neurons located in the CCe, LCa, and EG.

**Figure 3.**
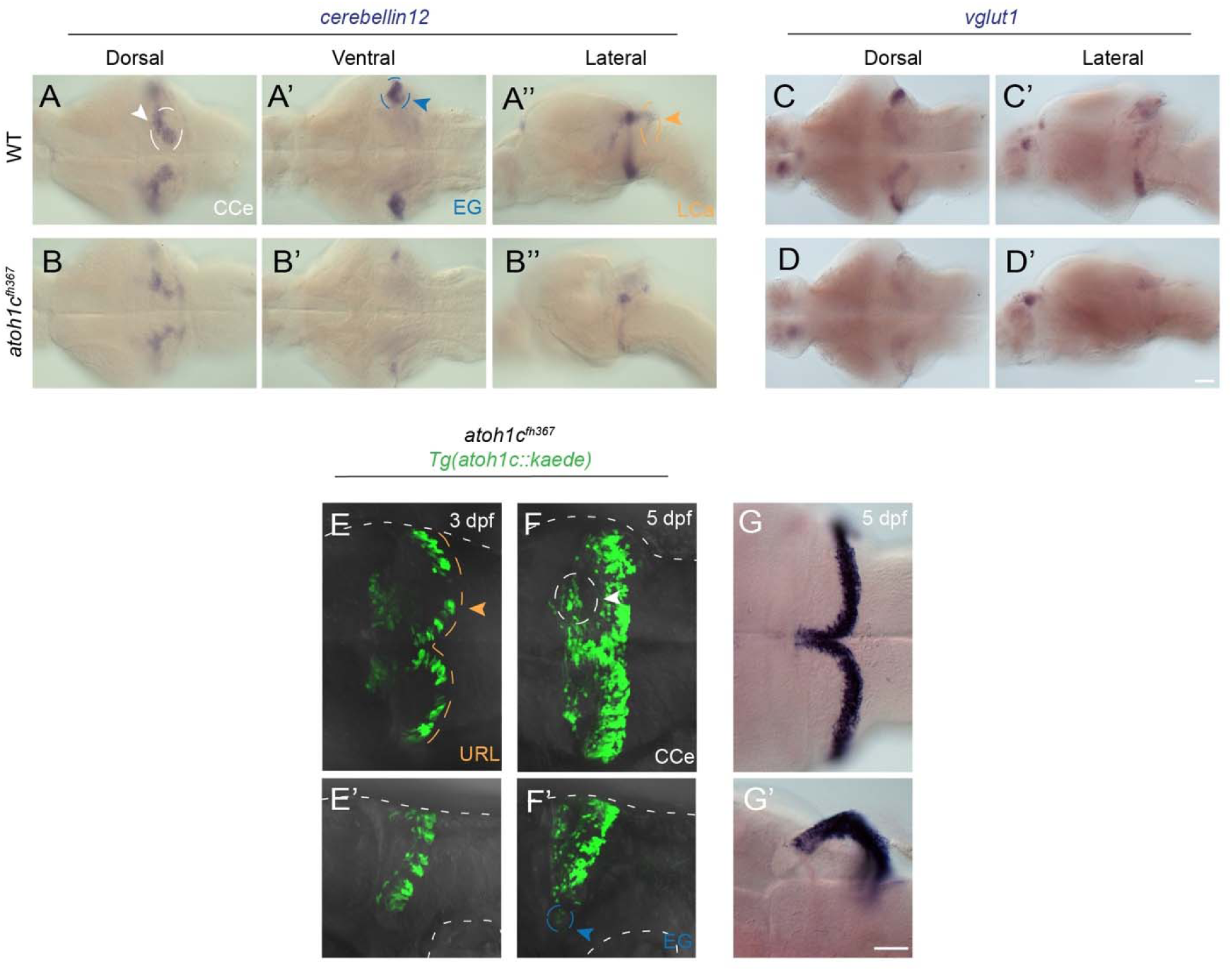
atoh1c is required for cerebellar granule neuron fate. A-D: Reduction in the number of mature granule neurons in atoh1c mutant as indicated by decreased staining for *cerebellin12* (B) and *vglut1* (D), at 5 dpf. E-G: The majority of mutant cells expressing the *Tg(atoh1c::kaede)* remain as an non-migrated population in the URL at 3 dpf (E compare to wild-type in Fig. 2 G,I) and 5 dpf (F compare to wild-type in Fig. 2J,L). G: Upregulation of *atoh1c* mRNA in URL at 5 dpf in atoh1c mutant embryos (compare to Fig. 1M,N). Dorsal (A,B,C,D,E,F,G), ventral (A’, B’) and lateral (A”, B”,C’,D’,E’,F’,G’) views are shown with anterior to the left. Scale bars: 50 μM.

If *atoh1c* is required for the development of granule neurons, then we predict one or more aspects of granule neuron maturation are defective in the mutant. Using markers of post-mitotic neurons (HuC/D; (Kim et al., 1996)), committed granule neuron precursors (NeuroD1; (Kani et al., 2010)) and cell proliferation (EdU incorporation) we determined the differentiation state of the *atoh1c^fh367^* cells (Fig. 4). It is important to note that due to epigenetic silencing of the UAS element (Akitake et al., 2011), not all of the atoh1c-derived granule neurons are detectable in a given fish using the *atoh1c::kaede* transgene. In 5 dpf wild-type larvae, the majority of *Tg(atoh1c::kaede)*+ cells have migrated away from their birthplace at the URL although a small population of HuC/D+ (Fig. 4A), NeuroD1+ (Fig. 4C) granule neurons remain near the URL in the LCa where, together with granule neurons in the EG, they contribute to the parallel fibers that project to the cerebellar-like structure in the dorsal hindbrain (Takeuchi et al., 2015). In contrast, in the *atoh1c* mutant, the majority of the *Tg(atoh1c::kaede)*+ cells remain at the URL but do not express HuC/D or elaborate axons (Fig. 4B), but they do express NeuroD1 (Fig. 4D) suggesting that they are committed but undifferentiated granule neuron precursors. To determine whether these are proliferating progenitors, we exposed 3 dpf larvae to 5-ethynyl-2’-deoxyuridine (EdU) continually for two days before fixing at 5 dpf (Fig 4E,F). Any cells that were proliferating during that period will incorporate EdU. In the *atoh1c* mutant, the massively expanded *Tg(atoh1c::kaede)*+ population in the URL is post-mitotic during this period (Fig. 4F). This post-mitotic but undifferentiated progenitor state with strongly increased *atoh1c* mRNA expression in *atoh1c* mutants (Fig. 3 I,J) is consistent with a classic proneural function for Atoh1c in the URL as has previously been proposed (Gazit et al., 2004; Machold et al., 2007; Millimaki et al., 2007).

**Figure 4.**
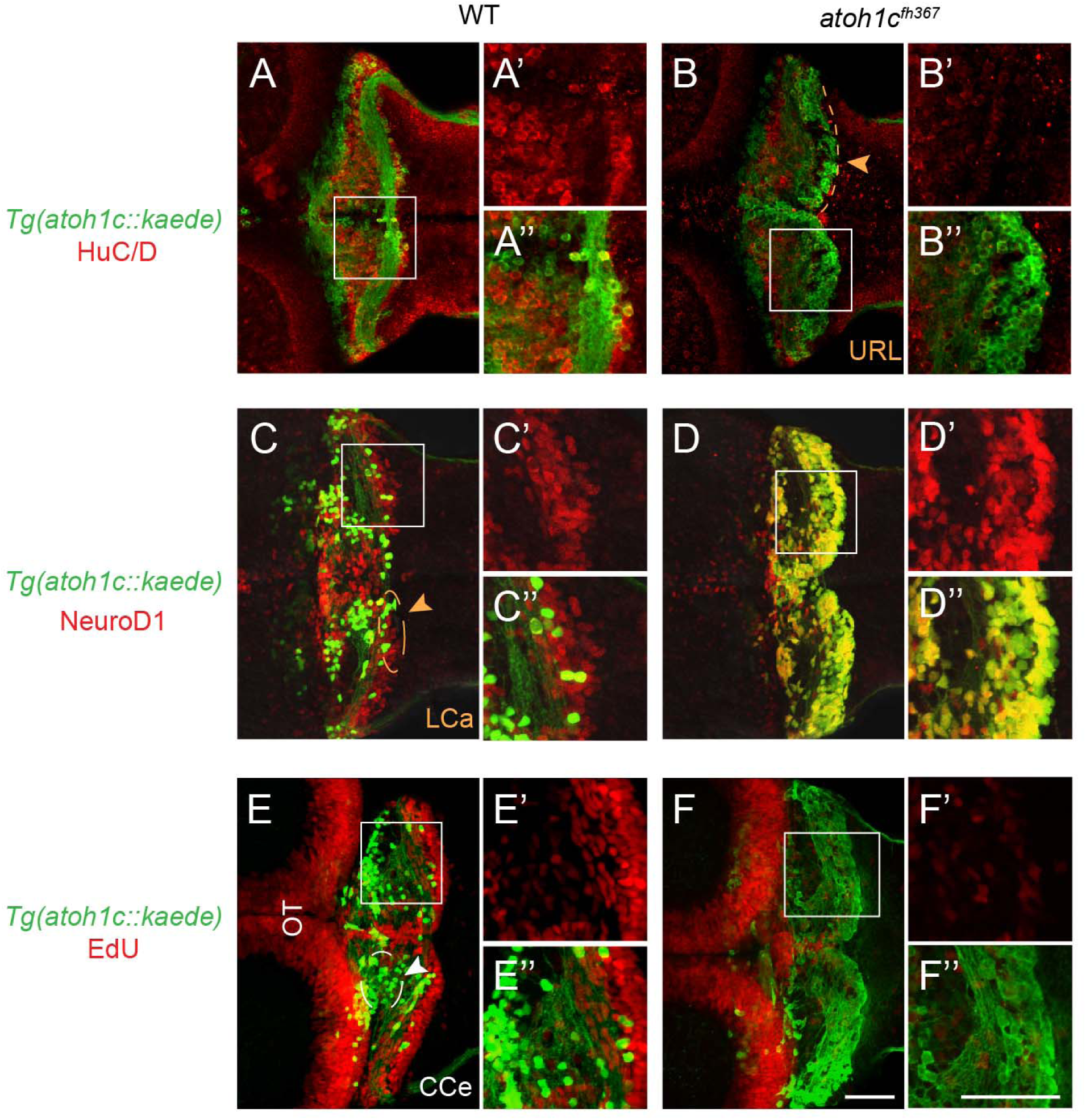
*atoh1c^fh367^* cerebellar cells accumulate as post-mitotic but undifferentiated granule progenitors. A,B: The majority of *Tg(atoh1c::kaede)*+ cells (green) in the *atoh1c* mutant are located within the URL and do not express HuC/D (red) at 5 dpf (B) indicating that they are not post-mitotic neurons. C,D: *atoh1c^fh367^* cells (green) in the URL are positive for NeuroD1 (red, D). E,F: *atoh1c^fh367^* cells (green) in the URL do not incorporate EdU (red, F). Strong EdU incorporation anterior to the cerebellum in both wild-type and mutant is in the optic tectum (OT). Dorsal views with anterior to the left. Scale bars: 50 μM.

### Atoh1c is required for delamination from the URL

Loss of *atoh1c* results in the accumulation atoh1c-expressing granule neuron precursors in the URL that are unable to terminally differentiate. How does a differentiation-arrested committed precursor behave? In order to address this, we performed high-resolution time-lapse imaging of both wild-type and *atoh1c^fh367^* embryos at 3 dpf, a time when we can continuously capture the birth and migration of granule neurons in wild-type embryos. Before migrating, wild-type *atoh1c*-expressing progenitors are epithelial, with their apical end feet along the ventricle-facing surface of the URL. They elaborate a basal process and very soon thereafter they release their apical contact and move away from the URL following this process (Koster and Fraser, 2001; Volkmann et al., 2010). This transition is accomplished in a period of six hours (Fig. 5, Movie 3). In contrast, *atoh1c* mutant progenitors at the URL elaborate highly dynamic basal processes but fail to detach from the epithelium during the same period (Fig. 5E-H, Movie 4). Interestingly, “escaper” neurons in *atoh1c* mutants complete granule neuron differentiation and elaborate parallel fibers (Fig. 3F, cells in CCe). To determine if detachment from the URL is a cell-autonomous function of *atoh1c* we generated genetic chimeras. In chimeras, *atoh1c* mutant cells migrated equally poorly from the URL when surrounded by wild-type cells (48% cells migrated; n=422 cells, 10 embryos) as when surrounded by *atoh1c* mutant cells (56% cells migrated; n=122 cells, 21 embryos). Conversely, wild-type cells migrated equally well when surrounded by mutant cells (72% cells migrated; n=284 cells, 11 embryos) as by wild-type cells (77% cells migrated; n=53 cells, 15 embryos). Thus, Atoh1c promotes the release of the granule neuron precursors from the URL epithelium, an essential step in neuronal differentiation (Hartenstein et al., 1992; Pacary et al., 2012).

**Figure 5.**
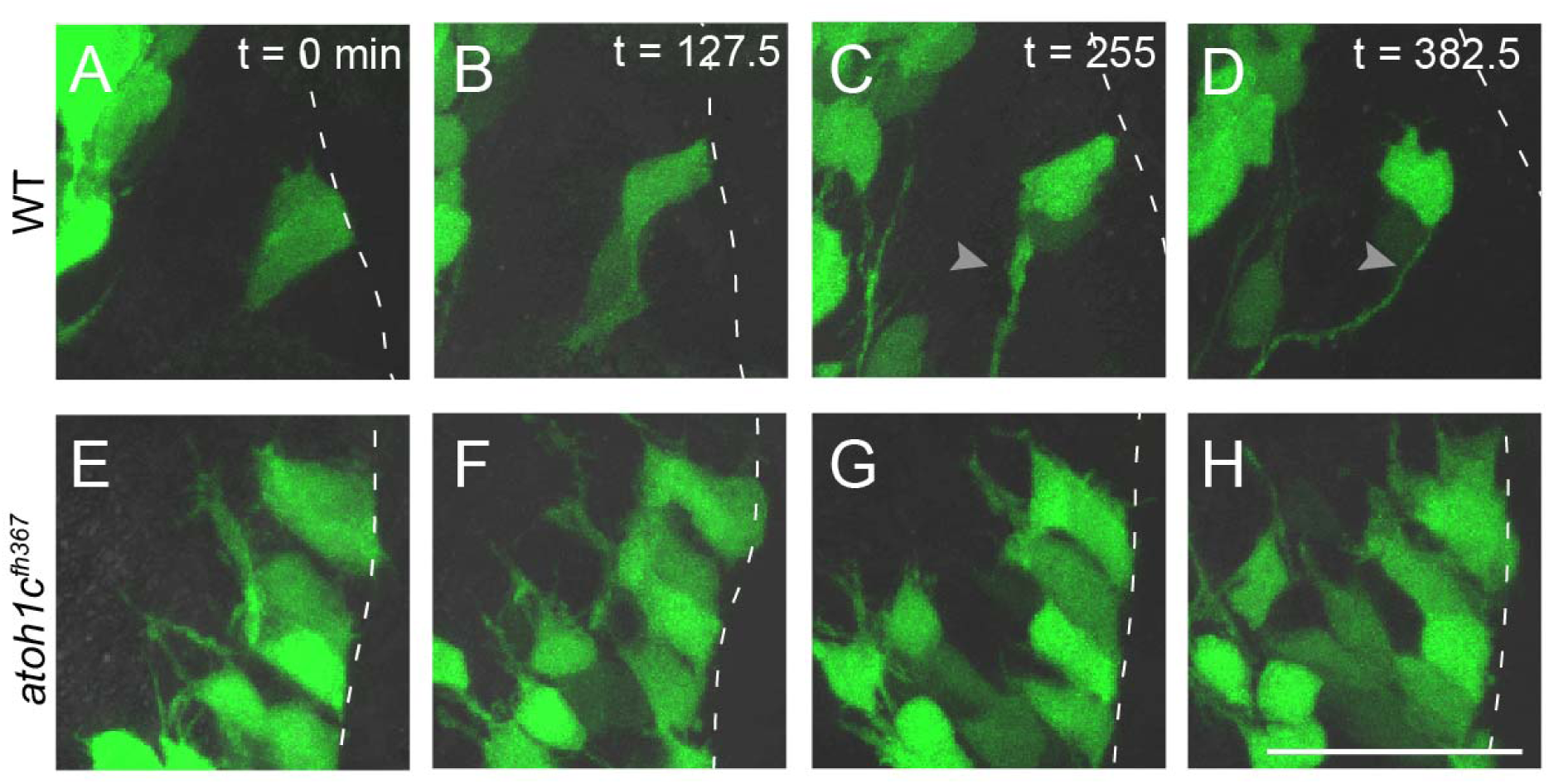
Atoh1c is required for release of granule neuron progenitors from URL. Still images from confocal timelapses taken at 3 dpf. Dotted line indicates the URL where the apical surfaces of granule neuron progenitors attach. In wild-type, *Tg(atoh1c::kaede)*+ progenitors detach from the URL over a six-hour interval (A-D) while in *atoh1c* mutants, *Tg(atoh1c::kaede)*+ cells remain attached at the URL (E-H). Time points indicated in upper right corner in minutes. Scale bar: 25 μM.

### Atoh1 paralogs function redundantly in granule neuron differentiation

Given their overlapping expression in the URL (Fig. 1), we were interested in understanding the functional relationships between the *atoh1* paralogues. We generated mutant alleles for *atoh1a* and *1b.* The *atoh1a^fh282^* allele is a missense mutation upstream of the predicted DNA binding domain and has been shown to have a loss-of-function phenotype in the zebrafish lateral line (Pujol-Marti et al., 2012). The *atoh1b^fh473^* allele is a 55 bp deletion that truncates the protein upstream of the predicted DNA binding domain. *atoh1a* and *atoh1c* are expressed independently of each other whereas the expression of *atoh1b* is regulated by *atoh1c* in regions of overlapping expression: the URL and midline of the CCe (not shown). Loss of function of *atoh1a* and *atoh1b* alone has no detectable effect on *cbln12* expression and the migration and differentiation of *Tg(atoh1c::kaede)*+ cells (Fig. 6B,C). In addition to its expression in the URL, *atoh1a* is expressed throughout the hindbrain lower rhombic lip (LRL) (Fig. S2A) which gives rise to a range of commissural interneurons and pre-cerebellar nuclei in mice (Bermingham et al., 2001; Helms and Johnson, 1998; Rose et al., 2009). Using an *atoh1a* reporter *(Tg(atoh1a:EGFP)nns7)* (Kani et al., 2010), we found that many *atoh1a:EGFP+* cells are retained at the LRL with a neural progenitor-like morphology in *atoh1a^fh282^* mutants (Fig. S2) suggesting a role for *atoh1a* in commissural interneuron differentiation as has been described in the mouse (Bermingham et al., 2001; Helms and Johnson, 1998; Rose et al., 2009).

**Figure 6.**
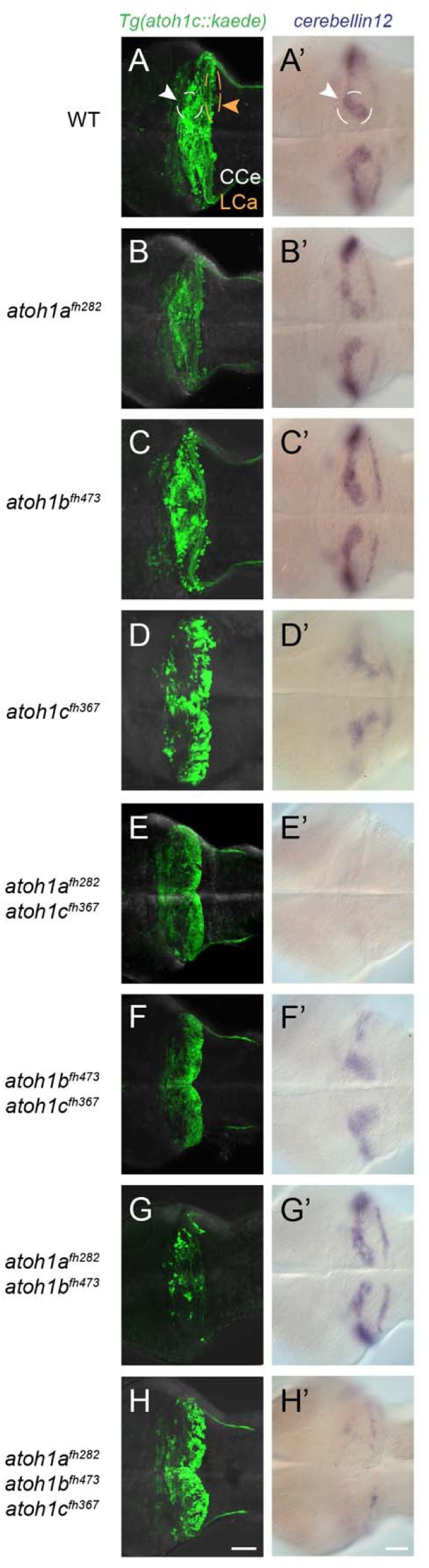
*atoh1a*, *atoh1b*, and *atoh1c* have non-overlapping roles in cerebellar development. Images of 5 dpf wild-type (A), single (B-D), and compound (E-H) *atoh1* mutants with the *Tg(atoh1c:kaede)* (left column) or *cbln12* RNA *in situ* hybridization (right column). Dorsal views with anterior to the left. Scale bars: 50 uM.

Although we observe a large reduction of mature granule neurons in the *atoh1c* mutant, some residual *cbln12* and *vglut1* expression remains, particularly in the CCe (Fig. 3B,D). To test possible redundancy amongst *atoh1* genes in granule neuron differentiation, we generated all possible *atoh1*^-/-^ allelic combinations. We found that in *atoh1a^fh282^*; *atoh1c^fh367^* double mutant embryos there was a near-complete loss of *cbln12* expression (Fig. 6E’). This was not further enhanced in the *atoh1a^fh282^; atoh1c^fh367^; atoh1b^fh473^* triple mutant (Fig. 6H’) suggesting that *atoh1b* does not contribute detectably to granule neuron development. We conclude that *atoh1a* has a minor role in granule neuron differentiation in the CCe that is only detectable in the absence of *atoh1c.* This is consistent with previous lineage analysis that showed only a minority of granule neurons in the CCe are derived from *atoh1a* progenitors (Kani et al., 2010).

### *atoh1a* and *atoh1c* have distinct derivatives in the developing cerebellum

The enhanced granule neuron phenotype in *atoh1a^fh282^; atoh1c^fh367^* double mutants could be either because *atoh1a* expression partially compensates for loss of *atoh1c* in the same progenitor cells, or because *atoh1a* and *atoh1c* are independently required for the differentiation of two different populations of granule neurons in the CCe, both of which express the pan-granule neuron marker *cbln12.* Consistent with the latter model, we detected no overlap in *Tg(atoh1c::kaede)* and *Tg(atoh1a:dtomato)*-expressing cells in double transgenic fish (Fig. 7) as well as by double RNA *in situ* hybridization (not shown). We conclude that the *Tg(atoh1a:EGFP)*+ granule neurons within the CCe described by Kani et al. (Kani et al., 2010) represent a population of *cbln12*+ granule neurons distinct from the majority *Tg(atoh1c::kaede)*+ population. It remains to be seen whether these *atoh1a*+ and *atoh1c*+ granule neuron populations do indeed play functionally distinct roles within the cerebellum.

**Figure 7.**
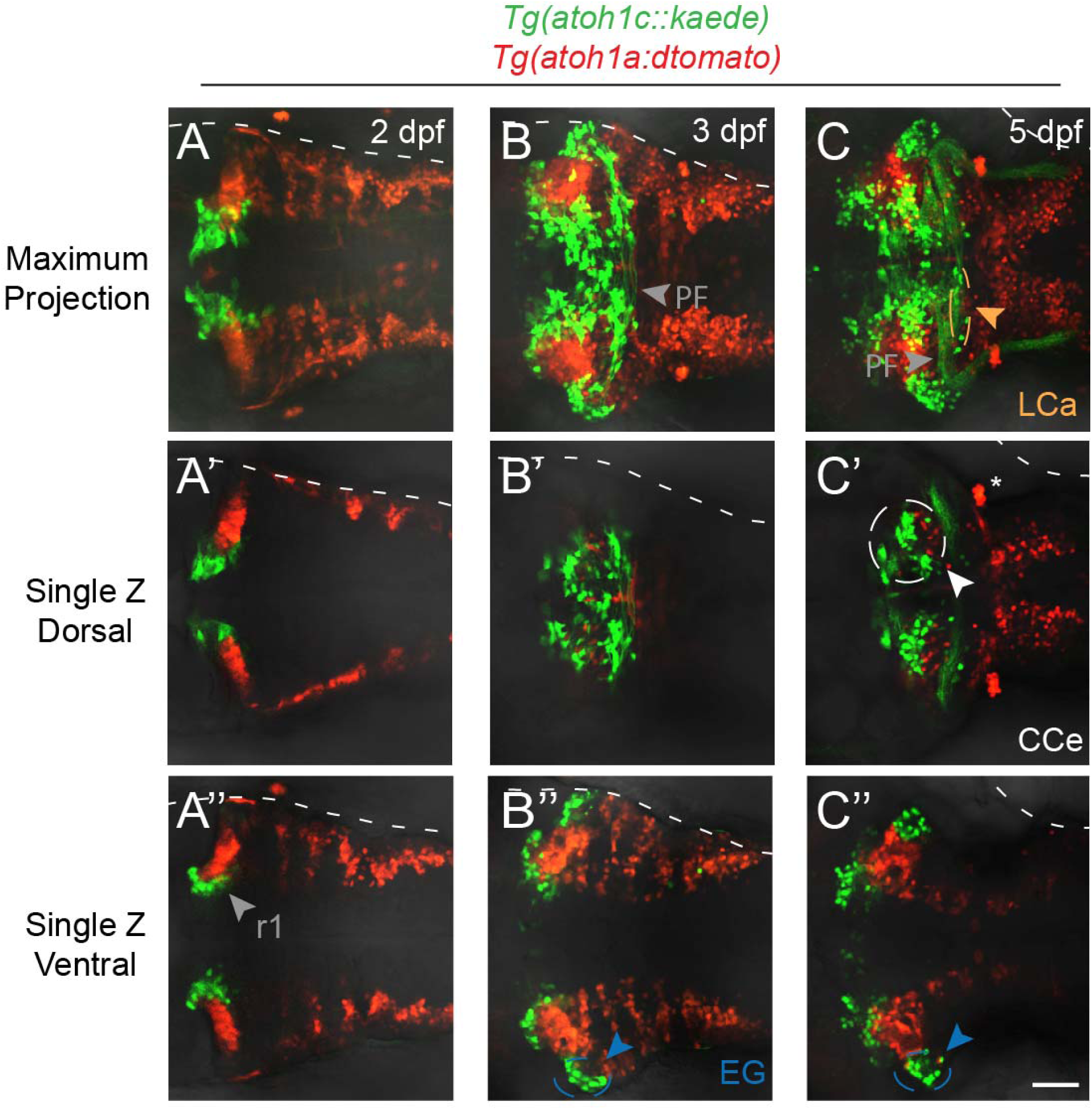
*atoh1a-* and *atoh1c-derived* neurons specify distinct progenitor pools. Live imaging of wild-type embryos with *Tg(atoh1c::kaede)* (green) and *Tg(atoh1a:dtomato)* (red) transgenes indicate *atoh1a* and atoh1c-derived neurons are distinct cerebellar populations. Maximum projections of a z-stack (A-C) and single z-slices at dorsal (A’-C’) or ventral (A”-C”) focal planes with anterior to the left. A: 2 dpf, B: 3 dpf, C: 5 dpf. Scale bar: 50 μM.

### *atoh1a* functionally rescues the *atoh1c^fh367^* phenotype

A classic model for the maintenance of duplicated genes posits that regulatory changes render both genes essential for a subset of their pre-duplicated functions (Force et al., 1999). If this is the case for *atoh1a* and *atoh1c*, we predict that either *atoh1a* or *atoh1c* could rescue granule neuron differentiation when expressed in the *atoh1c* progenitor domain in an *atoh1c^fh367^* mutant. We sought to rescue the *atoh1c^fh367^* phenotype by injection of a DNA construct encoding the full-length *atoh1a* cDNA under *atoh1c* regulation. We injected homozygous *atoh1c* mutant embryos carrying the *atoh1c:gal4ff* transgene at the one-cell stage with an *UAS:atoh1a-t2a-NLSmCherry* construct or a positive (*UAS:atoh1c-t2a-NLSmCherry*) or negative (*UAS:NLSmApple*) control construct, and determined the position of RFP-expressing cells at 4 dpf, using migration away from the URL as a proxy for the granule neuron differentiation. In these experiments, we used a very strict definition of “unmigrated” due to the nuclear localization of the RFP, scoring any cells with labeled nuclei that were not at the ventricular surface of the URL as “migrated”. We thus likely underestimated the proportion of unmigrated neurons. Nevertheless, we found that *atoh1a* (95% cells migrated; n=42 cells, 19 embryos) and *atoh1c* (96% cells migrated; n=27 cells, 11 embryos) were equally effective in rescuing the migration of *atoh1c* mutant granule neurons, compared to *UAS:NLSmApple* alone (80% cells migrated; n=172 cells, 17 embryos) We conclude that *atoh1a* and *atoh1c* have equivalent functions in granule neuron progenitors, at least with respect to inducing delamination from the URL.

## DISCUSSION

Here we have defined the roles of *atoh1* genes in zebrafish cerebellar development with the use of live imaging of transgenic reporters combined with mutant analysis. We find that of the three *atoh1* genes in the zebrafish genome, *atoh1c* plays the most prominent role in the development of cerebellar granule neurons, both in the cerebellar corpus (CCe) required for postural control and in the caudolateral lobes (EG and LCa) involved in vestibular function. We show that *atoh1c* is required not for granule neuron specification per se, but for granule neuron progenitors to lose their epithelial character, migrate away from the upper rhombic lip, and terminally differentiate. Finally, we show that *atoh1c* functions as an Wnt and FGF “coincidence detector” at the mid-hindbrain boundary to specify an early population of neuronal progenitors that give rise to a ventral r1 neurons related to the Locus Coeruleus.

### More atoh1 genes for more granule neuron diversity

Lineage tracing experiments in mouse have shown that *atoh1*-expressing progenitors at the URL give rise sequentially to tegmental, deep cerebellar and cerebellar granule neurons (Fig. 8) (Machold and Fishell, 2005; Rose et al., 2009; Wang et al., 2005). In this study, we have used the conserved cerebellar circuitry of the zebrafish to understand the role of the *atoh1* genes in specification of cerebellar neuronal populations. We consider granule neurons first. Based on *atoh1* expression patterns and lineage tracing of *atoh1a*+ cells in zebrafish which showed the presence of atoh1a-derived granule neurons in the Va and CCe but not the EG or LCa, Kani et al. predicted that *atoh1c and/or atoh1b* would be required for the granule neurons in these structures. Our work confirms this prediction: zebrafish *atoh1c* is required for the differentiation of EG and LCa granule neurons. Thus, additional *atoh1* genes during teleost evolution may have allowed for the formation of cerebellar structures such as the EG and the LCA that are not observed in mammals.

**Figure 8.**
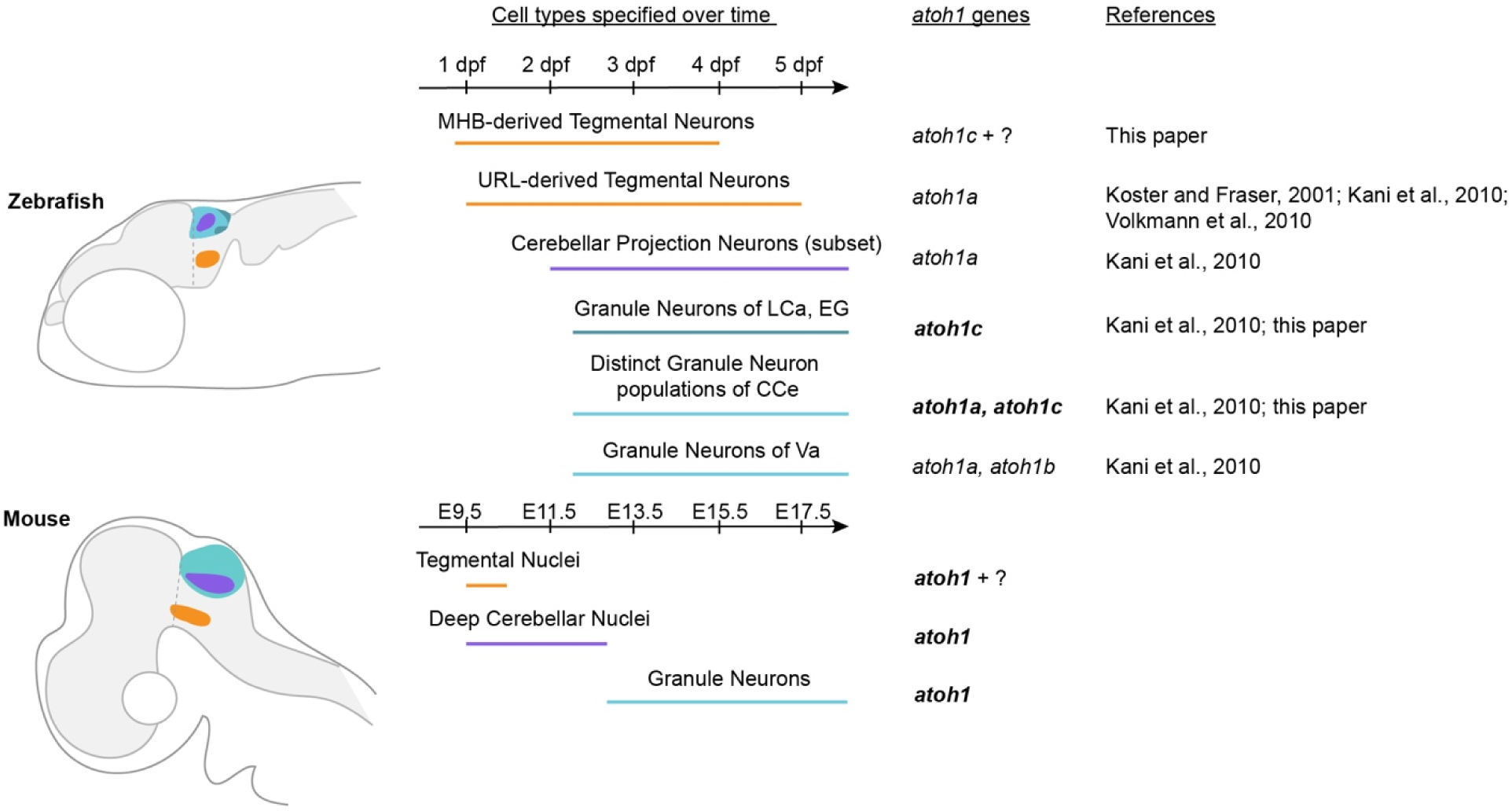
*atoh1*-derivatives in zebrafish and mouse. Schematic outlines *atoh1*-derivatives generated over time in the anterior hindbrain of zebrafish (top) and mouse (bottom). Bold text indicates gene has been shown to be required for development of the neuronal population. Homologous neuronal populations are indicated with the colored lines. Dpf: days post fertilization; E: embryonic day; CCe, corpus cerebelli; EG, eminentia granularis; LCa, lobus caudalis cerebelli; Va, valvula cerebelli.

Unexpectedly, our work reveals an additional degree of granule neuron diversity within the CCe, a homologous structure to the mammalian cerebellar vermis. We find that, *atoh1c* is also required for a population of granule neurons in the CCe that does not overlap with the *atoh1a* population, so that *atoh1c* and *atoh1a* together are required for the full complement of granule neurons in the CCe. Cerebellar granule neurons have been described as one singular population in vertebrates; however, our data reveals distinct *atoh1a* and *atoh1c* populations within the CCe (Fig. 8). Both populations express the pan-granule cell marker *cerebellin12*, as do the EG and the LCa, and our overexpression experiment suggests that *atoh1a* can substitute for *atoh1c* in the atoh1c-derived population. While it is possible that the two populations function identically in the cerebellar circuit, in a recent RNA-Seq screen for granule neuron-specific genes, Takeuchi et al. (Takeuchi et al., 2016) identified several granule neuron markers that have distinct patterns of expression within the CCe, suggesting heterogeneity in that population. It remains to be determined whether these domains correspond to the *atoh1a*-derived and *atoh1c*-derived granule neurons and whether the two populations are functionally distinct within the cerebellar circuit.

### atoh1 genes and the neuroepithelial progenitor state

We have observed that in the absence of *atoh1c*, progenitors fail to migrate away from the URL and differentiate into granule neurons. Instead, *Tg(atoh1c::kaede)* cells accumulate at the URL in a post-mitotic, partially differentiated (NeuroD1-positive, HuC/D-negative) state. A similar accumulation of granule neuron progenitors at the URL was detected in mouse *atoh1* mutants (Ben-Arie et al., 1997; Ben-Arie et al., 2000). Our finding that non-migrated *atoh1c* mutant progenitors at the URL express NeuroD1 is interesting in light of the fact that in mice, granule neurons do not turn on NeuroD1 until after they complete their transit-amplifying divisions in the external granule cell layer (Miyata et al., 1999). The expression of NeuroD1 in undifferentiated granule neuron progenitors at the URL in *atoh1c* mutants may reflect the finding that most granule neurons in fish differentiate directly after leaving the URL, without going through a transit-amplifying phase (Butts et al., 2014; Chaplin et al., 2010).

The accumulation of *atoh1c+* cells is typical of classical Notch-dependent lateral inhibition, whereby proneural genes limit their own expression by transcriptional activation of Delta, which in turn stimulates Notch in adjacent cells to inhibit proneural gene expression (Artavanis-Tsakonas et al., 1999). Our high-resolution live imaging of these cells in *atoh1c* mutants shows that they exhibit a migratory behavior, extending dynamic basal processes similar to wild-type granule progenitors do. However, while wild-type granule progenitors are able to release their apical contacts and migrate away from the URL, *atoh1c* mutant cells cannot release these contacts and remain trapped at the URL. Recent studies have shown that eliminating N-cadherin-containing apical junctions is a critical step in neuronal differentiation (Matsuda et al., 2016; Pacary et al., 2012; Rousso et al., 2012). Interestingly, *atoh1c* mutant progenitors that escape the URL appear to complete differentiation and generate parallel fibers. We also noted that the morphology of the *atoh1a^fh282^* cells in the LRL (Fig. S2B) resemble that of *atoh1c^fh367^* cells at the URL, suggesting that a key function of *atoh1* genes may be to promote apical detachment of neural progenitors, possibly by directly or indirectly repressing N-cadherin expression or activity.

### atoh1c-expressing cells at the MHB boundary give rise to tegmental neurons associated with the Locus Coeruleus

We have identified a novel population of atoh1c-derived neurons closely associated with the Locus Coeruleus, an evolutionarily ancient noradrenergic population involved in states of arousal and sleep-wake cycles. Like the LC neurons, the *atoh1c* neurons are specified immediately posterior to the MHB in cells that require the coincident reception of Wnt and FGF signals. They migrate ventrally and caudally into ventral r1, express tyrosine hydroxylase, and elaborate rostral, caudal, and cross-midline projections. However, unlike the LC, they do not express subsequent enzymes required for norepinephrine synthesis such as dopamine beta-hydroxylase (DBH, data not shown) and they turn off TH expression soon after arriving in ventral r1.

Lineage tracing in mouse has revealed an array of *atoh1*-derivatives including a number of tegmental pontine and deep cerebellar nuclei that appear before granule precursors emerge from the URL (Fig. 8) (Machold and Fishell, 2005; Rose et al., 2009; Wang et al., 2005). These include TH-positive cholinergic neurons in the lateral parabrachial and pedunculopontine tegmental nuclei, which lie adjacent to the Locus Coeruleus, and are vital for arousal and attention and have been implicated in the generation of REM sleep (Rose et al., 2009). In a separate study, early-born *atoh1-* derived neurons in the peri-LC region were shown to regulate sleep states in mice (Hayashi et al., 2015). We hypothesize that the MHB population of LC-associated *atoh1c* neurons may play an ancient role in regulating states of arousal, possibly analogous to the lateral parabrachial or pedunculopontine tegmental nuclei. However, because these *atoh1c*+ r1 neurons persist in all *atoh1* mutant combinations we have generated, we are as yet unable to assess their functions in mutant fish. Interestingly, Machold and Fishell also noted that some *atoh1*-derived tegmental neurons are present in *atoh1* mouse mutants, suggesting the presence of compensating mechanisms for the specification of some Atoh1-derived neuronal populations in mouse.

Green et al. (Green et al., 2014) recently identified an early *atoh1* expression domain at the mid-hindbrain boundary in chick and mouse which arises in response to MHB signals independently of the URL *atoh1* expression domain and gives rise to early-born *atoh1* neurons in the tegmentum. Given the early onset of *atoh1c* expression at the MHB in zebrafish and its dependence on MHB-derived Wnt1 and Fgf signaling, it is likely that this domain, and its tegmental derivatives, are homologous to the tetrapod MHB *atoh1* population described by Green et al.

### Concluding remarks

We have shown that *atoh1* genes contribute to the developmental of neuronal diversity in the zebrafish cerebellum, however we have not addressed their functions. Registering *atoh1*-derived populations with recent zebrafish neuroanatomical atlases will help to predict their functions (Marquart et al., 2015; Randlett et al., 2015), as will functional studies using genetically-encoded neuronal activity reporters under different behavioral paradigms(Muto and Kawakami, 2016). Surprisingly, *atoh1c* mutants are viable and do not exhibit obvious behavioral abnormalities as adults, however we anticipate that behaviors such as the vestibulo-ocular reflex, in which the eye rotates to compensate for changes in body position (Bianco et al., 2012), may be affected. Likewise, the hypothesized role of the *atoh1c*-expressing r1 neurons in arousal can be measured based on their activity in sleep paradigms (Chiu et al., 2016).

## MATERIAL AND METHODS

### Zebrafish Lines and Maintenance

Zebrafish (*Danio rerio*) were staged and maintained according to standard procedures as previously described (Kimmel et al., 1995). Experiments using zebrafish followed the Fred Hutchinson Cancer Research Center Institutional Animal Care and Use Committee standards and guidelines (IACUC#1392). All transgenic lines were maintained in the *AB background. Transgenic lines used in this study include *Tg(UAS:kaede)^s1999^* (Davison et al., 2007), gift of the Baier lab, *Tg(hsp70l:dnfgf1r-EGFP)^pd1^* (Lee et al., 2005), *Tg(hsp70:TCFΔC-GFP)^w74^* (Martin and Kimelman, 2012), gifts of the Kimelman lab, *Tg(atoh1a:EGFP)^nns7^* (Kani et al., 2010) and *Tg(atoh1a:dTomato)^nns8^* (Wada et al., 2010), gifts of the Hibi lab.

### Fgf and Wnt signaling inhibition

Fgf and Wnt signaling was blocked by incubating 10 hpf embryos containing the heat-inducible *Tg(hsp70l:dnfgfr-EGFP)^pd1^* or *Tg(hsp70:TCFΔC-GFP)^w74^* for 15 minutes at 38°C or 40°C, respectively (Lee et al., 2005; Martin and Kimelman, 2012).

### Generation of mutant alleles

*atoh1a^fh282^* was generated by TILLING (Draper et al., 2004) and has been previous described (Pujol-Marti et al., 2012). *atoh1b^fh473^* was generated with use of CRISPR/Cas9 as outlined in (Shah et al., 2015). The CRISPR gRNA sequences for *atoh1b* were: 5’-GGCTGACCCGGAGTGACCCG and 5’-GGTGCGTGCGTAATTCTCCA. *atoh1c^fh367^* was generated with the use of TALENs as outlined in (Sanjana et al., 2012). The TALEN target sequences for *atoh1c* were as follows: 5’-GGAGAGAGACTGACAGATC and 5’-CCAGCCCATTGGGGCTTT. In each experiment in this paper, embryos were phenotyped blind and then later genotyped by PCR using the following protocols: *atoh1a^fh282^:* forward primer 5’-ATGGATGGAATGAGCACGGA and reverse primer 5’-GTCGTTGTCAAAGGCTGGGA followed by digestion with AvaI (New England Biolabs) generates a 195 bp + 180 bp WT allele and a 195 bp + 258 bp mutant allele; *atoh1b^f^*: forward primer 5’-TGGACACTTTCGGGAGGAGT and reverse primer 5-CTTCAGAGGCAGCTTGAGGG generates a 180 bp WT allele and a 125 bp mutant allele; *atoh1c^fh367^:* forward primer 5’-GACTCCCTGTgGtCATTATCAA and reverse primer 5’-AGCTCACTCAGGGTGCTGAT generates a 510 bp WT allele and a 388 bp mutant allele.

### Plasmid construction and injection

*TgBAC(atoh1c:gal4ff)^fh430^* was generated according to the protocol outlined in (Bussmann and Schulte-Merker, 2011) and using BAC recombineering plasmids provided by the Shulte-Merker lab. A BAC containing *atoh1c* (#38G3) was obtained from the BACPAC Resources Center at Children’s Hospital Oakland Research Institute. This BAC was recombineered to insert *gal4ff* 114bp downstream of the *atoh1c* ATG and to add *tol2* arms flanking the genomic DNA insert. *Tg(UAS:kaede)^s1999^* embryos were injected at the one-cell stage with 180 pg of the modified BAC along with 90 pg of *tol2 transposase* mRNA. Expressing embryos were raised to adulthood for the establishment of stable transgenic lines.

### RNA in situ hybridization

For all *in situ* hybridizations, embryos were fixed in 4% paraformaldehyde with 1x PBS (phosphate-buffered saline) and 4% sucrose at 4°C overnight. RNA in situ hybridization was performed as described (Thisse et al., 1993). Embryos were mounted in glycerol on coverslips and transmitted light images were taken on a Zeiss Axioplan2 microscope.

### Live imaging

Embryos were anesthetized with 0.4% ethyl 3-aminobenzoate methanesulfonate (ms-222) (Sigma) and immobilized in 1.2% low-melting point agarose (Gibco). All embryos were imaged on a Zeiss LSM700 inverted confocal microscope.

### EdU Labeling

Starting at 3 dpf, embryos were incubated in final concentration of 0.5 mM F-ara-EdU (Sigma T511293) diluted in fish water. Embryos were anesthetized and fixed at 5 dpf. For EdU visualization, fixed whole embryos were permeablized in PBSTr (phosphate-buffered saline + 0.5% Triton X-100) for 30 mins at RT and then incubated in a solution containing 10uM Cy5-azide (Lumiprobe A2020), 2 mM copper(II) sulfate (Sigma 45167), and 20 mM sodium ascorbate (Sigma A7631) for 1 hour at RT. After 3 PBS washes, samples were processed for immunofluorescence as described below.

### Immunofluorescence

For whole-mount immunostaining, embryos were fixed in 4% paraformaldehyde with 1x PBS (phosphate-buffered saline) and 4% sucrose at 4°C overnight. The fixed embryos were washed with PBSTr (PBS + 0.5% Triton X-100), dissected and incubated in acetone at −20 °C for 7 min. Dissected brains were washed once with PBSTr and twice with PDT (PBS, 1% BSA, 1% DMSO, 0.5% Triton X-100), and incubated in 5% goat serum in PDT at RT for 2 h. The samples were incubated with the primary antibody solution at 4 °C overnight. After four washes with PBST, the tissues were incubated with secondary antibodies (1/250 dilution, Alexa Fluor 488 and/or Alexa Fluor 594 goat antimouse and/or goat anti-rabbit IgG (H + L), (Molecular Probes, Invitrogen). Following staining, tissue was cleared step-wise in a glycerol series and mounted for confocal imaging. The following antibodies were used: chicken anti-GFP (1:500, Abcam, ab13970); rabbit anti-Kaede (1:500, MBL Co. Ltd., PM012M); mouse anti-HuC/D (1:500, Invitrogen, A-21271); mouse anti-TH (1:500, Millipore, MAB318); mouse anti-NeuroD1(1:500, gift from the Hibi Lab).

### Chimeric Analysis

*atoh1c^fh367^, dnFgfr*, or *TCFΔC* embryos or wild-type controls to be used as donors in transplantation experiments were injected at the 1-cell stage with either Cascade Blue-dextran or Rhodamine-dextran (10,000 mw, Invitrogen). Cells were transplanted from these donor embryos into wild-type or *atoh1c^fh367^* mutant host embryos at the early gastrula stage as described (Kemp et al., 2009), targeting donor cells to the dorsal CNS at the mid-hindbrain level of host recipients. Donor embryos were raised to 3 dpf to determine which transgene(s) they carried, and/or were genotyped for the presence of the *atoh1c^fh367^* allele. Hosts of donors of the relevant genotype(s) were imaged as described above (“Live Imaging”).

## ACKNOWLEGDEMENTS

We thank David Kimelman, Rachel Wong, David Raible, Herwig Baier, and Stephan Schulte-Merker for transgenic lines, constructs and reagents. David Prober and Owen Randlett provided valuable help regarding the identity of *atoh1c* r1 neurons. We would like to thank the members of the Moens lab for helpful discussions and comments on this manuscript, Jason Stonick for technical help, and Rachel Garcia for excellent zebrafish care. We also thank Luyuan Pan, Arish Shah, and Daniel Berman for their help with the isolation of the *atoh1c^fh367^* mutant.

## COMPETING INTERESTS

The authors declare no competing financial interests.

## AUTHOR CONTRIBUTIONS

C.B.M. and C.U.K. designed and performed experiments, analyzed data, and wrote the manuscript. C.Y.S. performed experiments for supplemental figure 1. M.H. provided key reagents prior to publication as well as intellectual input into the interpretation of the data.

## FUNDING

This work was supported by National Institutes of Health R01 NS082567 to C.B.M. and National Institutes of Health Training Grant T32 HD007183 to C.U.K.

